# Photosynthetic-product-dependent Activation of Plasma Membrane H^+^-ATPase and Nitrate Uptake in Arabidopsis Leaves

**DOI:** 10.1101/2022.06.30.498222

**Authors:** Satoru N. Kinoshita, Takamasa Suzuki, Takatoshi Kiba, Hitoshi Sakakibara, Toshinori Kinoshita

## Abstract

Plasma membrane (PM) H^+^-ATPase is a pivotal enzyme for plant growth and development that acts as a primary transporter and is activated by phosphorylation of the penultimate residue, threonine, at the C-terminus. Photosynthetically active radiation activates PM H^+^-ATPase via phosphorylation in mesophyll cells of *Arabidopsis thaliana*, and phosphorylation of PM H^+^-ATPase depends on photosynthesis and photosynthesis-related sugar supplementation, such as sucrose, fructose and glucose. However, the molecular mechanism and the physiological role of photosynthesis-dependent PM H^+^-ATPase activation are still unknown. Analysis using sugar analogues, such as palatinose, turanose, and 2-deoxy glucose, revealed that sucrose metabolites and products of glycolysis such as pyruvate induce phosphorylation of PM H^+^-ATPase. Transcriptome analysis showed that novel isoform of the *Small Auxin Up RNA* genes, *SAUR30*, is upregulated in a light- and sucrose-dependent manner. Time course analyzes of sucrose supplementation showed that phosphorylation level of PM H^+^-ATPase increased within 10 min, but expression level of *SAUR30* increased later than 10 min. The results suggest two temporal regulations may participate in the regulation of PM H^+^-ATPase. Interestingly, a ^15^NO_3_^-^ uptake assay in leaves showed that light increases ^15^NO_3_^-^ uptake, and that increment of ^15^NO_3_^-^ uptake depends on PM H^+^-ATPase activity. The results opened the possibility of physiological role of photosynthesis-dependent PM H^+^-ATPase activation in the uptake of NO_3_^-^. We speculate that PM H^+^-ATPase may connect photosynthesis and nitrogen metabolism in leaves.

## Introduction

Photosynthesis in plants converts CO_2_ and light energy to carbon metabolites for translocation and energy production and is important for plant growth and reproduction. To maintain high photosynthetic activity, several inorganic nutrients and elements are essential, such as nitrogen, potassium, magnesium, and phosphorus (Tränkner et al., 2018; Evans and Clarke, 2019). The nitrogen content of leaves is of utmost importance for photosynthetic capacity. Most of nitrogen in C_3_ sun leaves is allocated to photosynthesis-related proteins such as light-harvesting complex, Rubisco, and enzymes in the Calvin–Benson cycle (Evans and Clarke, 2019). It is thus essential to maintain high nitrogen availability in leaves for active photosynthesis. To this end, vascular plants must uptake nitrate or other nitrogen metabolites from soil and translocate nitrogen to leaves in the form of nitrate or amino acids (Delhon et al., 1996; Matt et al., 2001).

Uptake of nitrogen—including as nitrate, ammonium, and amino acids—requires transport across the plasma membrane (PM). Therefore, nitrogen uptake is mediated by PM-localized transporters and channels. Among the nitrogen transporters and channels, nitrate transporters (NRTs) have been well-characterized in a range of plant species, and most NRTs, such as the NRT1 and NRT2 families, are H^+^-symporters (Hachiya and Sakakibara, 2017). The association of the H^+^ gradient across the PM with NRTs in roots has been investigated in several plant and crop species (Tong et al., 2005). The association of the H^+^ gradient across the PM with nitrate uptake in leaves of Arabidopsis and cucumber has been suggested (Cookson et al., 2005; Nikolic et al., 2012).

The H^+^ gradient and membrane potential are maintained and polarized by H^+^ extrusion from the cell, which is mediated by active H^+^ pumps, primarily PM H^+^-ATPase driven by ATP hydrolysis (Sondergaard et al., 2004). The physiological role of PM H^+^-ATPase in plants is not limited to nutrient uptake, also encompassing dormancy alleviation in seeds (de Bont et al., 2019), cell elongation in hypocotyl (Takahashi et al., 2012), stomatal opening in guard cells (Assmann et al., 1985; Kinoshita and Shimazaki, 1999), sugar loading in phloem (DeWitt and Sussman, 1995), and pollen tube growth in flower pollen (Robertson et al., 2004; Hoffmann et al., 2020). Double mutation of the two major isoforms of PM H^+^-ATPase (*aha1 aha2*) in *Arabidopsis thaliana* leads to embryo death (Haruta et al., 2010), implying that PM H^+^-ATPase is essential for plants and that revealing its function by genetic approaches will be difficult. The H^+^ pump activity of PM H^+^-ATPase is regulated not only by its abundance but also by post-translational modification (PTM) (Falhof et al., 2016). One of the major PTM regulators is phosphorylation of the penultimate residue, threonine (Thr), at the C-terminus. Phosphorylation of the penultimate Thr simultaneous with 14-3-3 protein binding to the region activates H^+^ pumping (Palmgren, 2001). Therefore, regulation of penultimate Thr phosphorylation by specific protein kinases and protein phosphatases determines the activity of PM H^+^-ATPase in plants.

The signaling and regulatory mechanisms of PM H^+^-ATPase have been identified. Phytohormones such as auxin and brassinosteroid induce phosphorylation of PM H^+^-ATPase via the *Small auxin up RNAs* (*SAURs*)–protein phosphatase 2C-D clade (PP2C-D) module in the hypocotyl of seedlings, resulting in cell elongation (Takahashi et al., 2012; Spartz et al., 2014; Ren et al., 2018; Minami et al., 2019; Wong et al., 2019). In guard cells, blue-light illumination induces phosphorylation of PM H^+^-ATPase via phototropin-BLUS1-BHP modules, leading to opening of stomata (Inoue and Kinoshita, 2017). Several *SAUR*–PP2C-D modules have been implicated in PM H^+^-ATPase activation in guard cells (Wong et al., 2021; Akiyama et al., 2022). However, the activation mechanism of PM H^+^-ATPase in photosynthetic tissues is unclear.

We have reported that photosynthetically active radiation of thalli of *Marchantia polymorpha*, protonemata of *Physcomitrella patens*, or leaves of vascular plants induced phosphorylation of PM H^+^-ATPase (Okumura et al., 2012a; Okumura et al., 2012b; Okumura et al., 2016; Harada et al., 2020). Although the mechanism is dependent on photosynthesis and its products (Okumura et al., 2016), the molecular mechanism and physiological role of PM H^+^-ATPase activation in photosynthetic tissues are unknown.

Here, we characterized the molecular mechanism of photosynthesis-dependent phosphorylation of the penultimate Thr of PM H^+^-ATPase and the physiological role of activated PM H^+^-ATPase in Arabidopsis leaves. First, sugar-analogue supplementation to leaves showed that the phosphorylation of PM H^+^-ATPase is induced by glycolysis and the downstream metabolites. Second, a transcriptome analysis and time course experiment implicated two temporal regulatory mechanism in the phosphorylation of PM H^+^-ATPase. Finally, an isotope-labelled nitrate uptake assay implied that PM H^+^-ATPase activation by photosynthesis may have positive role in nitrate uptake in leaves.

## Results

### Involvement of glycolysis and the downstream in PM H^+^-ATPase phosphorylation in leaves

Light-induced phosphorylation of the penultimate residue, Thr, of PM H^+^-ATPase, which is required for PM H^+^-ATPase activation, is likely mediated by endogenous photosynthetic products such as sucrose, glucose, fructose, glucose-1-phosphate (G1P), and glucose-6-phosphatate (G6P) in mesophyll cells of *Arabidopsis thaliana* (Okumura et al., 2016; Supplemental Figure S1). Because these activating metabolites are involved in sugar signaling such as sucrose and hexose signaling (Li and Sheen, 2016), we used sugar analogues to assess whether phosphorylation of PM H^+^-ATPase is dependent on some types of sugar signaling.

First, the effects of exogenous sucrose and the sucrose analogues palatinose and turanose were investigated. Palatinose and turanose are not metabolized in plants, but induce the sucrose-specific signaling pathway (Fernie et al., 2001; Sinha et al., 2002). Palatinose, turanose, sucrose, and mannitol were exogenously supplemented to pieces of leaves from dark-adapted Arabidopsis in the dark. The effects of sugars on the phosphorylation status of the penultimate Thr, of PM H^+^-ATPase were determined using an antibody against the phosphorylated penultimate Thr of PM H^+^-ATPase (anti-pThr; Hayashi et al., 2010). Palatinose and turanose had no effect on the phosphorylation level of PM H^+^-ATPase, whereas sucrose induced PM H^+^-ATPase phosphorylation (**Figure 1A, 1B**), implying that PM H^+^-ATPase phosphorylation is independent from the sucrose-specific signaling pathway, but requires sucrose metabolism.

**Figure 1.**
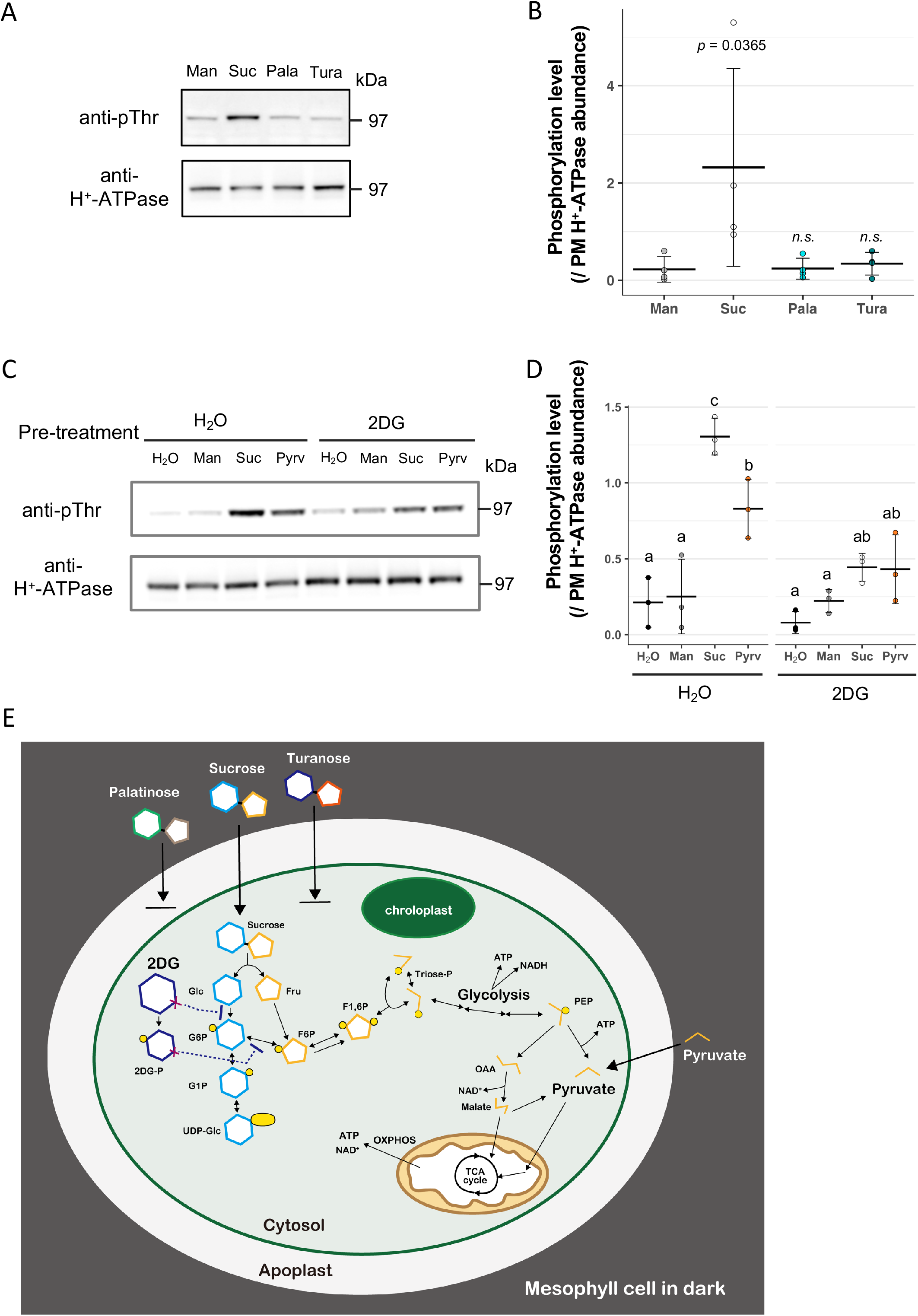
Glycolysis- and the downstream metabolites-induced PM H^+^-ATPase phosphorylation. A, Phosphorylation of the penultimate Thr and abundance of PM H^+^-ATPase in sucrose-analogue-supplemented pieces of leaves using anti-pThr and anti-PM H^+^-ATPase antibodies, respectively. Suc, Man, Pala, and Tura— sucrose, mannitol, palatinose, and turanose, respectively. B, Phosphorylation of PM H^+^-ATPase in sucrose-analogue-supplemented pieces of leaves. Cross bars and error bars represent means and SD of four independent experiments. *P-*values by one-way ANOVA with Dunnett’s test compared to the control; n.s., not significant (*P* > 0.05). C, Phosphorylation of penultimate Thr and abundance of PM H^+^-ATPase in metabolite-supplemented pieces of leaves using anti-pThr and anti-PM H^+^-ATPase antibodies. Pieces of leaves were preincubated with MilliQ water (H_2_O) or 2-deoxy glucose (2DG). Pyrv, pyruvate. D, Phosphorylation of PM H^+^-ATPase in metabolite-supplemented pieces of leaves. Cross bars and error bars represent means and SD of three independent experiments. Different letters above bars indicate significant differences by one-way ANOVA with Tukey HSD test (*P* < 0.05). E, Schematic of related metabolites and expected effects of sugar analogues. Exogeneous sucrose metabolism in dark is shown. The unmetabolized sucrose analogues palatinose and turanose are not converted to hexoses intracellularly. 2-deoxyglucose (2DG) is phosphorylated to 2DG-phosphate and competitively inhibits sucrose metabolism. Glc, glucose; Fru, fructose; G6P, glucose-6-phosphate; G1P, glucose-1-phosphate; F6P, fructose-6-phosphate; UDP-Glc, UDP-glucose; suc-6-P, sucrose-6-phosphate; F1,6P, fructose-1,6-bisphosphate; Triose-P, Triose-phosphate; PEP, phosphoenolpyruvate; OAA, oxaloacetate; OXPHOS, oxidative phosphorylation

Second, 2 deoxy-glucose (2DG) was used to investigate whether the phosphorylation of PM H^+^-ATPase requires hexose-phosphate accumulation and hexose signaling. 2DG is phosphorylated by hexokinases in cell but is not converted to fructose-6-phosphatae (F6P); thus, 2DG-phosphate accumulates intracellularly as hexose-phosphate (Klein and Stitt, 1998; Li and Sheen, 2016). In addition, 2DG inhibits carbon flux to the glycolysis pathway in animal and plant species (Xiong et al., 2013; Pajak et al., 2020). Pre-incubation of 2DG alone and with mannitol did not alter the phosphorylation level of PM H^+^-ATPase compared to the control (H_2_O alone), indicating that hexose-phosphate accumulation nor hexose signaling are not involved in the pathway (**Figure 1C, 1D**). Surprisingly, supplementation of pyruvate, the final product of glycolysis, induced phosphorylation of PM H^+^-ATPase. In addition, pre-incubation with 2DG suppressed sucrose-induced phosphorylation of PM H^+^-ATPase, whereas pyruvate-induced phosphorylation was less affected (**Figure 1C, 1D**). These results imply that metabolites of glycolysis and the downstream may be important for phosphorylation of PM H^+^-ATPase (**Figure 1E**).

### An activation mechanism of PM H^+^-ATPase via transcription of SAUR30

To unravel the molecular mechanism by which photosynthesis controls phosphorylation of PM H^+^-ATPase, the involvement of *SAUR* family genes was investigated. SAUR proteins activate PM H^+^-ATPase by inhibiting PP2C-D family proteins, which directly dephosphorylate penultimate Thr of PM H^+^-ATPase, in the hypocotyl of seedlings (Spartz et al., 2014; Ren et al., 2018; Wong et al., 2019). For this purpose, RNA-seq was performed to assess the expression levels of 79 *SAUR* family genes using dark-adapted Arabidopsis leaves subjected to white-light illumination (Lt), or incubation with 30 mM sucrose solution (Suc) for 30 min. Reference control samples for the Lt and Suc conditions were dark-adapted leaves (Dk) and 30 mM mannitol-solution-incubated leaves (Man), respectively.

Eighteen *SAUR* family genes were identified as differentially expressed genes (DEGs; FDR < 0.05, fold-change > 2) for the Lt/Dk comparison and four *SAUR* family genes for the Suc/Man comparison (**Figure 2**). Only *SAUR30* (AT5G53590) was differentially expressed in both Lt and Suc with high transcript per million (TPM) values (**Figure 2**). The expression of *SAUR30* was increased 3.9-fold in the Lt/Dk comparison and 2.3-fold in the Suc/Man comparison. Other than *SAUR* genes, 2,663 DEGs were detected in the Lt/Dk comparison, and 367 in the Suc/Man comparison (**Supplemental Figure S2A; Supplemental Data File 1**).

**Figure 2.**
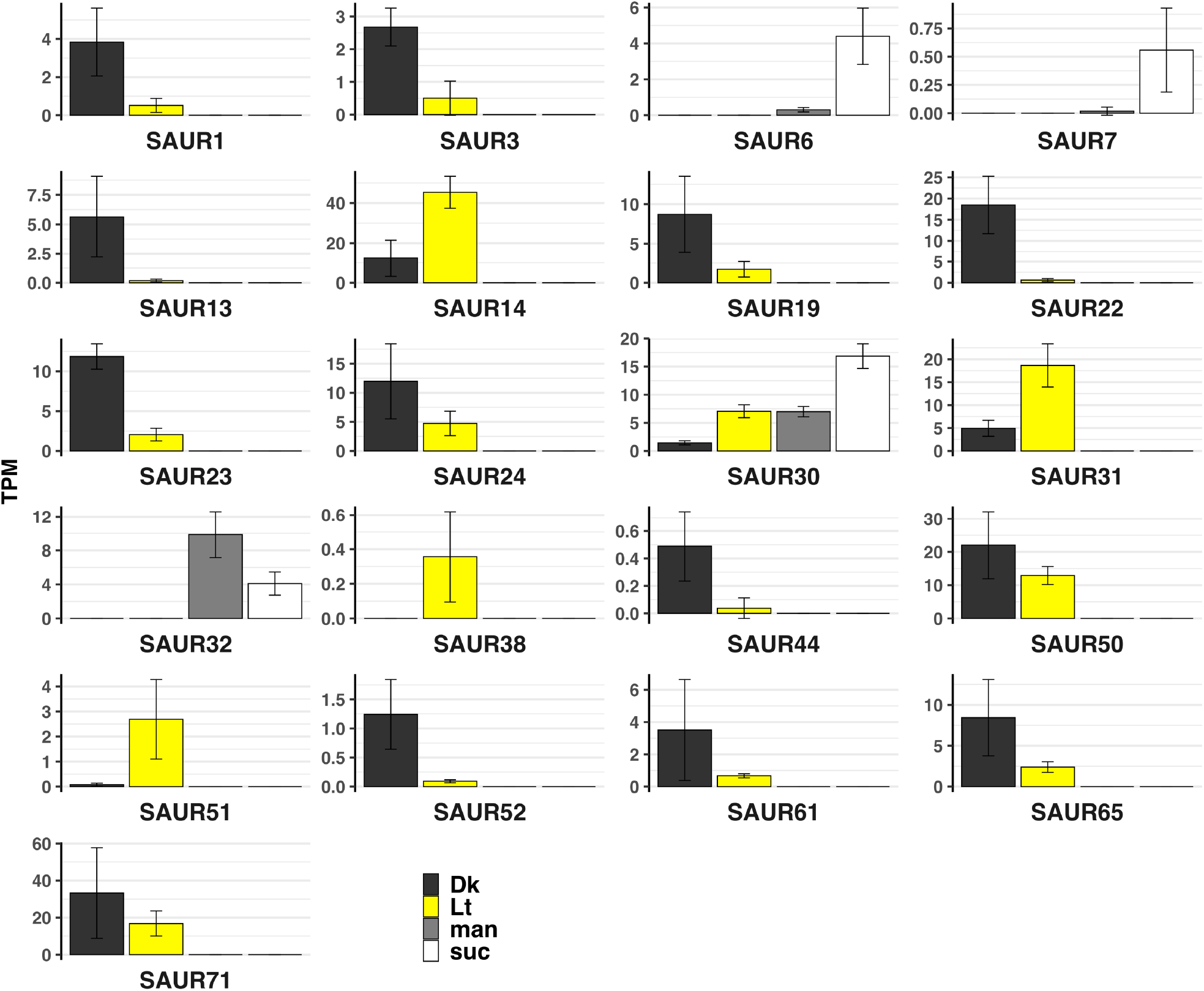
Differentially expressed *SAUR* genes in leaves. Transcripts per million (TPM) values of SAUR genes differentially expressed under Light or Sucrose condition. Dk, Lt, Man, and Suc—dark, light, mannitol, and sucrose, respectively. Bars and error bars represent the means and SD of four independent experiments.

Additionally, 23% (62/270) of DEGs under both conditions overlapped with KIN10-dependent reference DEGs (Baena-González et al., 2007) (**Supplemental Figure S2A**). KIN10 is a kinase subunit of SnRK1, a hub regulator of energy starvation. The overlapping genes showed reversed expression changes compared to those caused by KIN10 overexpression (Baena-González et al., 2007) (**Supplemental Figure S2B**), indicating that the leaf samples in our experiment were experiencing energy starvation after dark-adaptation, and energy recovery under light illumination and with exogenous sucrose supplementation (**Supplemental Figure S2C**).

Next, to confirm the function of SAUR30 in PM H^+^-ATPase phosphorylation *in vivo*, a *SAUR30-* containing plasmid was transfected into mesophyll cell protoplasts (MCPs) using the polyethylene glycol (PEG)-Ca^2+^ method, and phosphorylation of endogenous PM H^+^-ATPase in MCPs was examined. As expected, transiently expressed SAUR30 and SAUR30-GFP proteins increased the phosphorylation level of PM H^+^-ATPase in MCPs, whereas GFP alone and the no-protein-expression control (NC) did not (**Figure 3; Supplemental Figure S3**) We further confirmed that the high phosphorylation level of PM H^+^-ATPase in SAUR30-expressed MCPs was decreased when co-expressed with mutated PP2C-Ds (**Supplemental Figure S4**). The results indicate that the novel isoform of *SAUR, SAUR30*, may maintain PM H^+^-ATPase phosphorylation in MCPs via well-known SAUR-PP2C-D module model (Wong et al., 2019).

**Figure 3.**
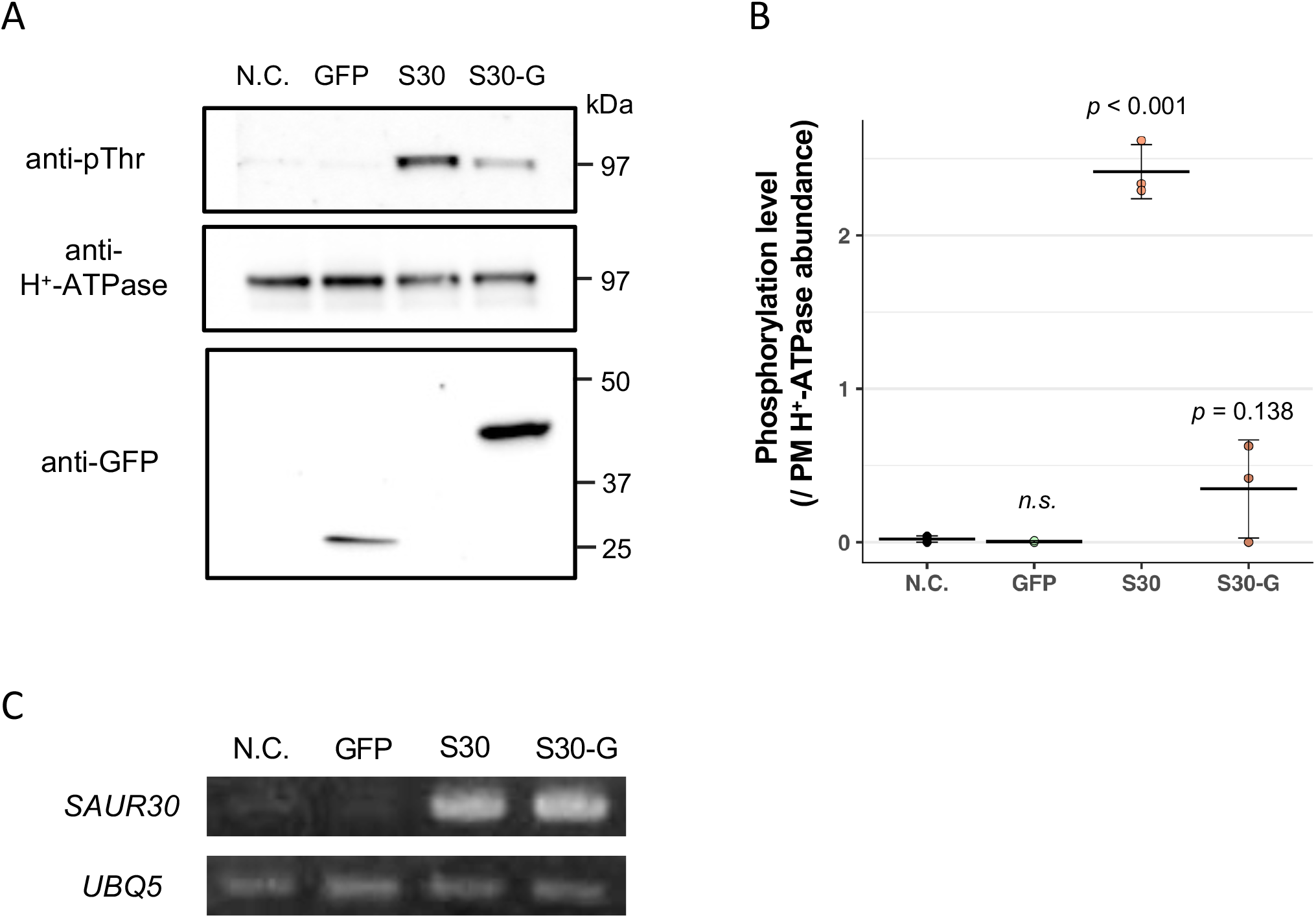
SAUR30-induced PM H^+^-ATPase phosphorylation in mesophyll cell protoplasts. A, Phosphorylation of the penultimate Thr and abundance of PM H^+^-ATPase in MCPs. The band at ∼ 25 kDa is GFP and that at 37–50 kDa is SAUR30-GFP. B, Phosphorylation of PM H^+^-ATPase in transfected MCPs. Cross bars and error bars represent means and SD of three independent experiments. *P-*values by one-way ANOVA with Dunnet’s test compared to the control; n.s., not significant (*P* > 0.05). C, Detection of *SAUR30* overexpression in transfected MCPs by RT-PCR. *UBQ5* was used as the internal control.

### Earlier regulatory mechanisms of PM H^+^-ATPase phosphorylation

The light-dependent phosphorylation of PM H^+^-ATPase starts 15 min after white light illumination (Okumura et al., 2016). The quick response of PM H^+^-ATPase to photosynthesis implies that phosphorylation of PM H^+^-ATPase is not only regulated by transcription in leaves. To examine the temporal pattern of PM H^+^-ATPase phosphorylation and *SAUR30* transcription in sucrose supplementation, pieces of leaves infiltrated with sucrose solution were sampled at 0, 2.5, 5.0, 10, 30, 60 min after infiltration. Next, the phosphorylation level of PM H^+^-ATPase and the expression level of *SAUR30* were evaluated by immunoblotting and RT-quantitative PCR, respectively. Interestingly, PM H^+^-ATPase phosphorylation was induced at around 5–10 min (**Figure 4A, 4B**), while *SAUR30* expression was induced markedly at 30 min after sucrose infiltration (**Figure 4C**). These results implicate another regulatory mechanism (5–10 min; early response) in phosphorylation of PM H^+^-ATPase in leaves.

**Figure 4.**
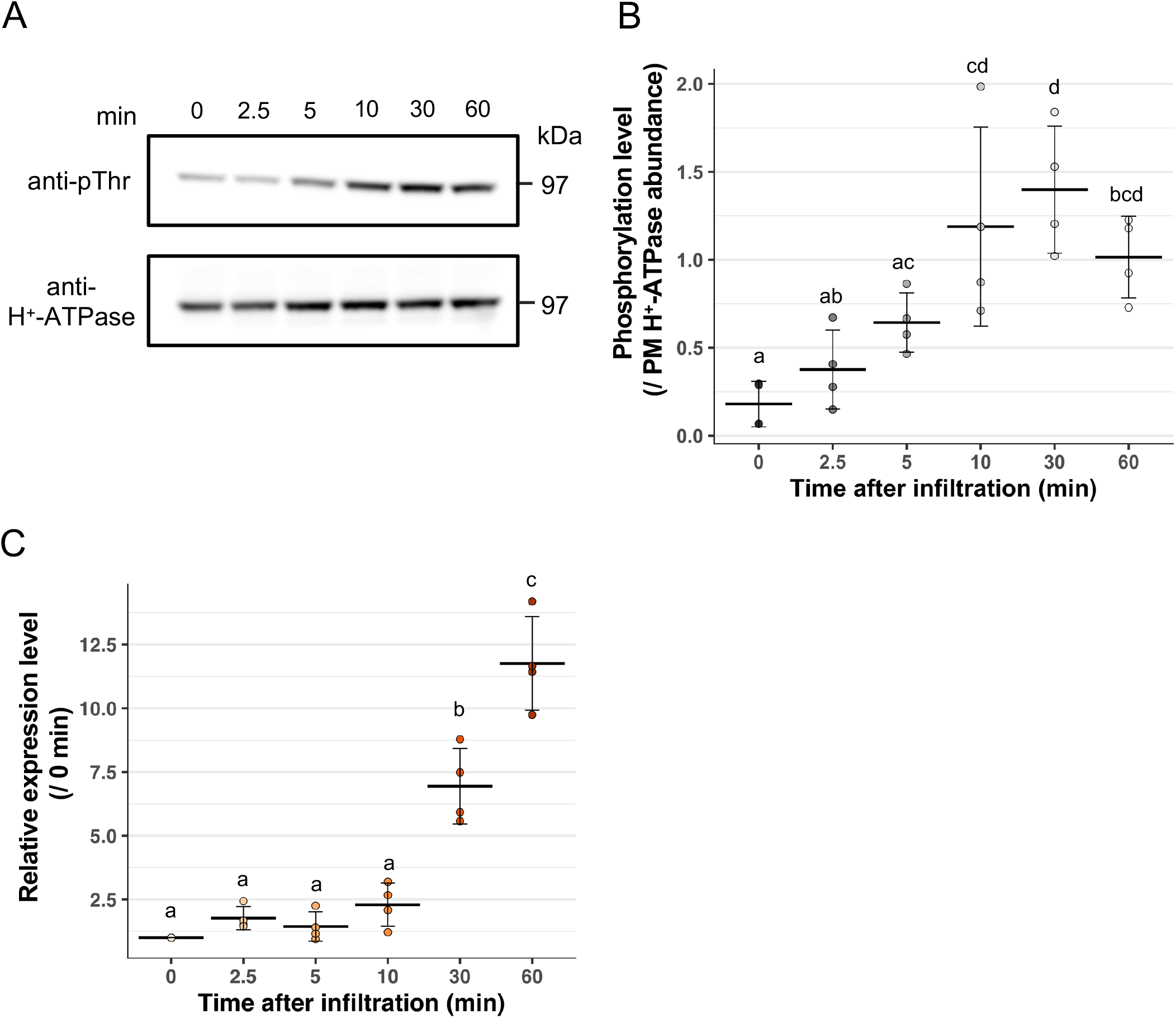
Temporal difference between phosphorylation of PM H^+^-ATPase and expression of *SAUR30* in sucrose-supplemented leaves. A, Phosphorylation of the penultimate Thr and abundance of PM H^+^-ATPase in pieces of leaves supplemented with sucrose as determined using anti-pThr and anti-PM H^+^-ATPase antibodies. B, Phosphorylation of PM H^+^-ATPase in pieces of leaves supplemented with sucrose. Pieces of leaves were flash frozen at the indicated times after infiltration. Cross bars and error bars represent the means and SD of four independent experiments. Different letters above bars indicate significant differences by one-way ANOVA with Tukey HSD test (*P* < 0.05). C, Expression of *SAUR30* in pieces of leaves supplemented with sucrose. Pieces of leaves were flash frozen at the indicated times after infiltration. *UBQ5* was used as the internal control. Cross bars and error bars represent the means and SD of four independent experiments. Different letters above bars indicate significant differences by one-way ANOVA with Tukey HSD test (*P* < 0.05).

### Positive role of PM H^+^-ATPase in light-dependent nitrate uptake

In illuminated leaves, nitrogen assimilation is triggered by the activation of nitrate reductase (NR) in response to photosynthesis (Kaiser and Huber, 2001). In addition, PM H^+^-ATPase generates an H^+^ gradient across the PM, which supports several H^+^ gradient-dependent transporters, including PM-localized nitrate transporters (Hachiya and Sakakibara, 2017). In line with the phosphorylation level of PM H^+^-ATPase, ATP hydrolysis activity of PM H^+^-ATPase in illuminated leaves increases compared to dark condition (Okumura et al., 2016). Therefore, we hypothesized that light-induced PM H^+^-ATPase activation in leaves acidifies apoplasts to generate an H^+^ gradient and supports nitrate transporters.

To test the above hypothesis, the nitrate uptake of leaves with different PM H^+^-ATPase activities was investigated using 1 mM K^15^NO_3_ (**Figure 5A**). As expected, illuminated leaves showed increased nitrate uptake compared to dark-adapted leaves. Moreover, the fungal toxin fusicoccin (Fc), an irreversible activator of PM H^+^-ATPase, significantly increased nitrate uptake in dark, whereas vanadate, an inhibitor of PM H^+^-ATPase (Palmgren, 2001), suppressed the increment of nitrate uptake under illumination (**Figure 5B**). Although the relationship between PM H^+^-ATPase activation and nitrate uptake need to be further investigated, these results imply the PM H^+^-ATPase activation may have a positive role in light-dependent nitrate uptake in leaves.

**Figure 5.**
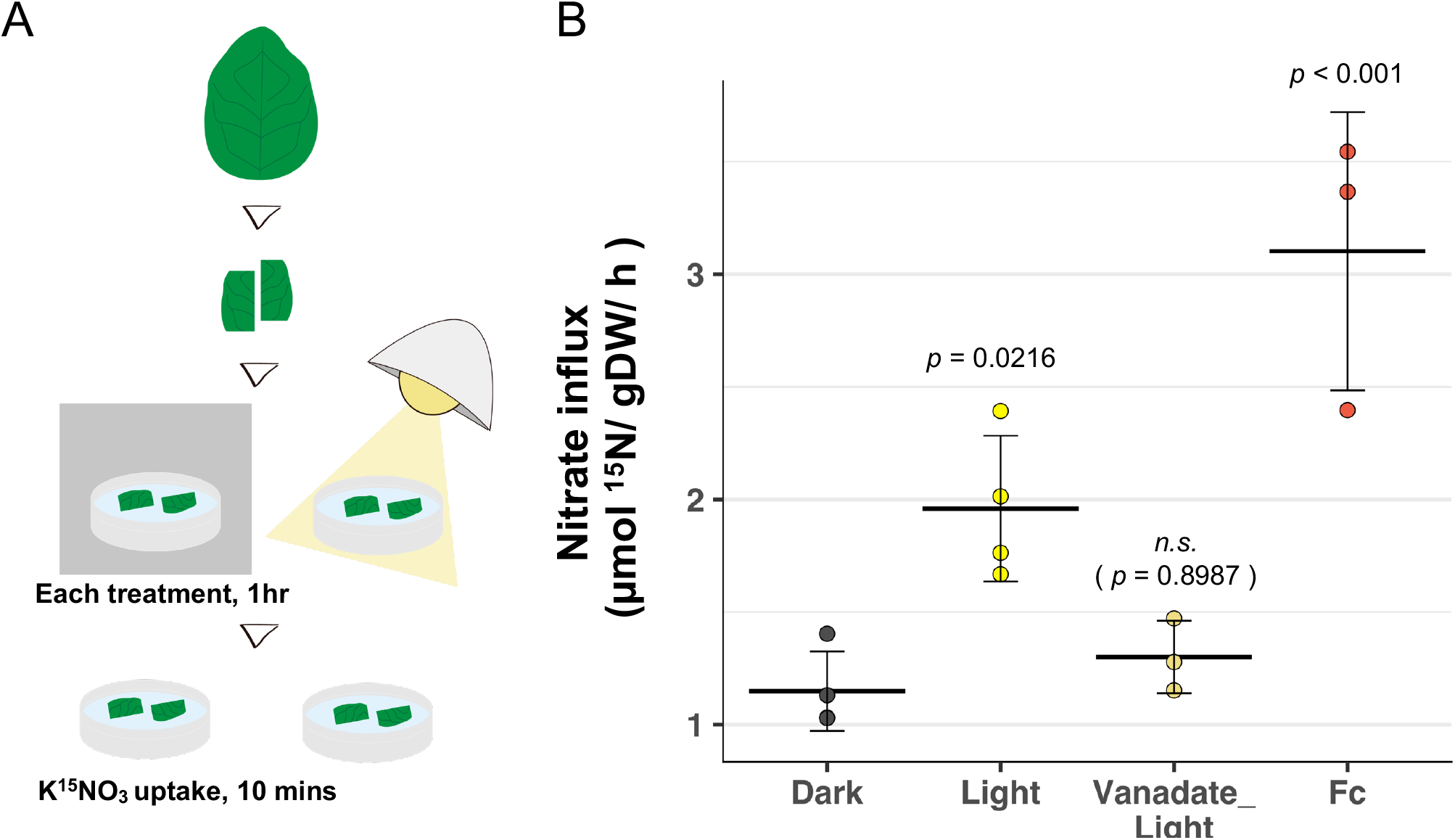
Light- and PM H^+^-ATPase-dependent nitrate uptake in leaves. A, Schematic of nitrate uptake assay. Leaves were harvested and cut in half, removing the main vein, and floated on water (Dark, Light), fusicoccin (Fc), and 1 mM vanadate in 10 mM Tris-HCl (pH 5.7) (vanadate_Light). For the vanadate_Light sample, pieces of leaves were preincubated with vanadate solution for 30 min and transferred to light conditions. After treatment, pieces of leaves were transferred to 1 mM K^15^NO_3_ solution and incubated for 10 min. B, Nitrate influx in leaves treated with Dark, Light, and a PM H^+^-ATPase inhibitor followed by Light (Vanadate_Light) or activator (Fc). Cross bars and error bars represent the means and SD of three to four independent experiments. *P-*values by one-way ANOVA with Dunnett’s test compared to the dark sample as the control; n.s. not significant (*P* > 0.05).

## Discussion

Photosynthesis- and photosynthetic-product-dependent phosphorylation of PM H^+^-ATPase has been reported in thalli of *Marchantia polymorpha* (Okumura et al., 2012a), protonemata of *Physcomitrella patens* (Okumura et al., 2012b), the leaves of *Arabidopsis thaliana* and several vascular plant species (Okumura et al., 2016), and leaves of *Vallisneria gigantea* (Harada et al., 2020). Sucrose-dependent phosphorylation of PM H^+^-ATPase was also reported in Arabidopsis seedlings (Niittylä et al., 2007). Although photosynthesis- and photosynthetic-product-dependent phosphorylation of PM H^+^-ATPase is conserved among a wide range of plant species, the molecular mechanism and physiological roles of the phosphorylation are unclear. Therefore, we used Arabidopsis leaves to evaluate the molecular mechanism and physiological role of PM H^+^-ATPase phosphorylation.

Our findings provide insight into the molecular mechanism and physiological role of PM H^+^-ATPase activation in leaves. First, photosynthetic-products, especially of glycolysis and the downstream, are important for inducing PM H^+^-ATPase phosphorylation. Second, a novel isoform of SAUR genes, *SAUR30*, is implicated in the activation mechanism via SAUR-PP2C-D modules, but an as-yet-unknown early regulatory mechanism was also implied. Finally, we suggest that PM H^+^-ATPase activation via photosynthesis may have a positive role in light-dependent nitrate uptake in leaves (**Figure 6**).

**Figure 6.**
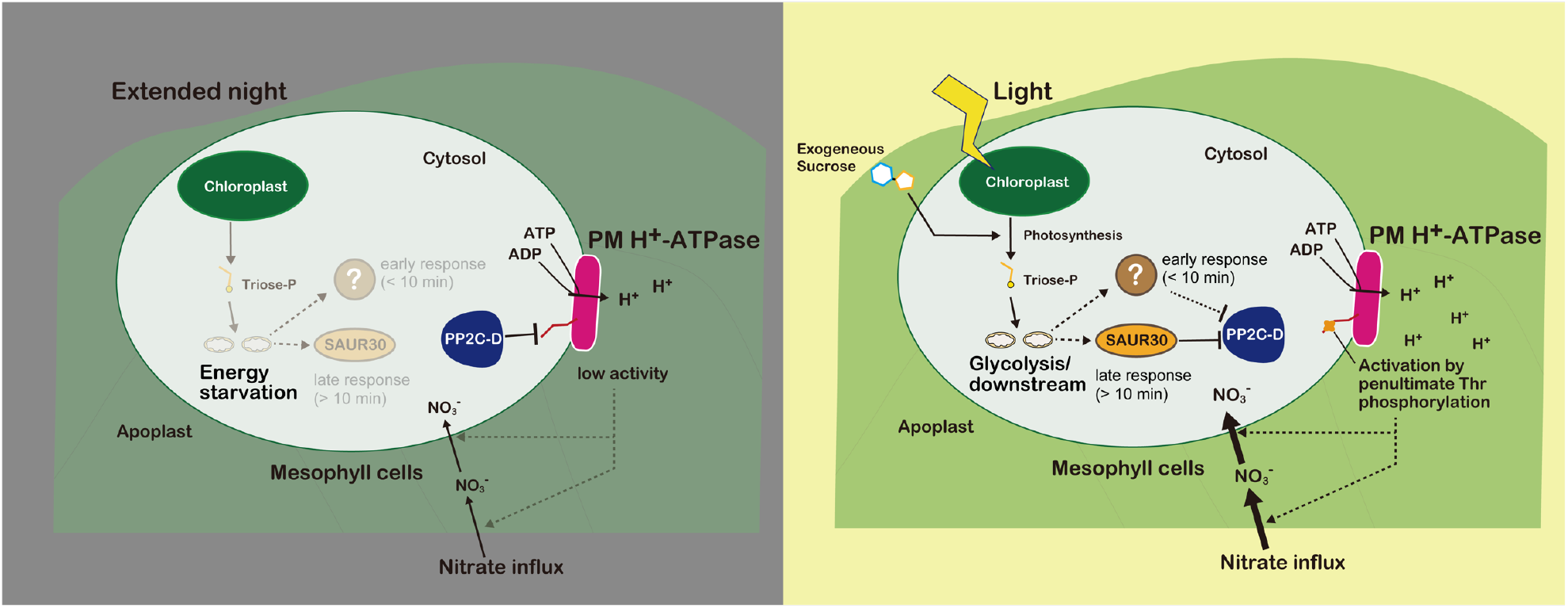
Hypothetical model of the mechanisms and physiological roles of PM H^+^-ATPase activation in leaves. Photosynthesis generates triosep-phosphate in illuminated leaves and glycolysis and the downstream metabolites induce the unknown activation mechanism within 5–10 min (early response) and SAUR30 expression after 10 min (late response). The inductions inhibit PP2C-D, leading to high phosphorylation level of PM H^+^-ATPase. Activated PM H^+^-ATPase creates an H^+^ gradient across the PM. The H^+^ gradient may facilitate nitrate uptake into leaves.

### Glycolysis and the downstream metabolite may participate in photosynthesis-dependent PM H^+^-ATPas phosphorylation in leaves

Sucrose is predominantly produced in source leaves for translocation to other sink tissues and is responsible for disaccharide signaling in plant cells (Li and Sheen, 2016). Palatinose and turanose are not metabolized in plants but induce the sucrose-specific signaling pathway (Fernie et al., 2001; Sinha et al., 2002). Palatinose is not recognized by the AtSUC2 transporter, a possible sucrose signal receptor, whereas turanose is recognized by the AtSUC2 transporter (Chandran et al., 2003). Only sucrose induced phosphorylation of PM H^+^-ATPase (**Figure 1A, 1B**). Therefore, sucrose-specific signaling pathway is not involved in PM H^+^-ATPase phosphorylation.

Next, involvement of hexose and hexose-phosphate signaling was investigated. Intracellular hexose availability is perceived by hexokinase (Li and Sheen, 2016). To mimic the intracellular accumulation of hexose and hexose-phosphate, the glucose analogue 2DG was applied to leaves. 2DG is phosphorylated intracellularly and perceived by hexokinase (Klein and Stitt, 1998; Li and Sheen, 2016). Supplementation of 2DG alone had no effect (**Figure 1C, 1D**), indicating that PM H^+^-ATPase phosphorylation is independent of hexose accumulation and the hexose-phosphate pool.

In addition, 2DG inhibits glycolysis in cancer cells (Pajak et al., 2020) and plant seedlings (Xiong et al., 2013). 2DG-pretreated leaves showed decreased PM H^+^-ATPase phosphorylation with exogenous supplementation of sucrose, whereas exogenous supplementation of pyruvate had a lesser effect with 2DG (**Figure 1C, 1D**). Therefore, respiration carbon metabolites may be responsible for phosphorylation of PM H^+^-ATPase, but not sugar production. This is consistent with the transcriptome analysis results implying that dark-adapted plants recovered from energy starvation after light illumination or sucrose supplementation (**Supplementary Figure S2**). However, this is tentative, because the carbon flux in illuminated leaves differs from that of leaves in darkness.

In illuminated leaves, mitochondrial pyruvate dehydrogenase (PDH; AT1G53240), a rate-limiting enzyme of pyruvate conversion to Acetyl-CoA in mitochondria and the entry point to the tricarboxylic acid (TCA) cycle, is inhibited by transcriptional and post-translational modification (Zhang et al., 2021). Therefore, the TCA cycle is limited due to the inactivation of PDH in illuminated leaves, and the rate of oxidative phosphorylation is low (Sweetlove et al., 2010). Isotope-labeled carbon flux analysis revealed that several photosynthesis, glycolysis, and non-cyclic TCA reaction metabolites are produced within 20 min of illumination—2-phosphoglyceric acid, pyruvate, alanine, serine, and trehalose (Szecowka et al., 2013). Together with the previous report that PM H^+^-ATPase phosphorylation is induced after 15 to 30 min of illumination (Okumura et al., 2016), the finding that 2DG-pretreatent suppressed sucrose-dependent phosphorylation of PM H^+^-ATPase in the dark (**Figure 1C, 1D**) implicates glycolysis and the downstream metabolites in illuminated leaves in photosynthesis-dependent phosphorylation of PM H^+^-ATPase. Studies using inducible mutants that can transiently alter the specific related metabolic pathway may provide further insight.

### Two temporal mechanism of PM H^+^-ATPase in Arabidopsis leaves

Several *SAUR* genes activate PM H^+^-ATPase by inhibiting the protein phosphatase PP2C D-clade (PP2C-D; Ren et al., 2018). Although many *SAUR* genes are responsive to auxin, *SAUR30* is less responsive than other family members (Paponov et al., 2008). We found that *SAUR30* is likely responsive to photosynthetic products and was the most abundant *SAUR* in sucrose-supplemented leaves (**Figure 2**). Furthermore, transient expression of *SAUR30* in MCPs induced phosphorylation of endogenous PM H^+^-ATPase via inhibiting PP2C-Ds (**Figure 3; Supplemental Figure S4**), suggesting photosynthesis-dependent expression of *SAUR30* maintain high phosphorylation level of PM H^+^-ATPase in leaves. Other *SAUR* family genes, which were excluded from our analysis because of a high FDR value, may function redundantly with *SAUR30*. Other DEGs found in both light and sucrose condition may also participate in the pathway.

Notably, induction of *SAUR30* expression in sucrose-supplemented leaves occurred later than PM H^+^-ATPase phosphorylation (**Figure 4**), implying the involvement of an unknown early responsive mechanism (5–10 min) of PM H^+^-ATPase phosphorylation in leaves. We speculate that PTM of specific proteins, including SAUR30 proteins accumulated before illumination, or metabolite-protein interactions induce phosphorylation of PM H^+^-ATPase as soon as photosynthesis occurs, which is subsequently (> 10 min) supported by transcription of *SAUR30*. Further investigation of protein modification in leaves with sucrose supplementation may provide mechanistic insights.

### Possible role of PM H^+^-ATPase in light-dependent nitrate uptake of leaves

To dissect the physiological importance of PM H^+^-ATPase activation in leaves, we investigated the relationship between photosynthesis-dependent PM H^+^-ATPase activation and nitrogen assimilation in illuminated leaves, because photosynthesis and nitrogen assimilation are closely associated. Synthesis of major components of the photosynthesis machinery—including chlorophyll, light-harvesting complex, Rubisco, and ATP synthase—in chloroplasts requires large amounts of nitrogen (Evans and Clarke, 2019). In addition, the photosynthesis rate is significantly affected by nitrogen availability (Perchlik and Tegeder, 2018). PM H^+^-ATPase generates an H^+^ gradient across the PM. The activity of H^+^-coupled nutrient transporters is dependent on the PM potential and H^+^ gradient (Palmgren, 2001; Sondergaard et al., 2004). Therefore, we hypothesized that PM H^+^-ATPase activated by photosynthesis energize PM-localized nitrate transporters, such as NRT proteins. Although our stable isotope-labelled nitrate (^15^NO_3_^-^) uptake assay with pieces of leaves did not demonstrate direct evidence of involvement of NRT proteins nor actual cellular nitrate influx across PM, the nitrate uptake assay using cut piece of leaves implies that light-induced increment of nitrate uptake was dependent on PM H^+^-ATPase activity (**Figure 5**).

It has been suggested that transportation of nitrogen requires the PM H^+^-gradient and membrane potential in leaves of some plants including Arabidopsis and nitrogen-starved cucumber (Cookson et al., 2005; Nikolic et al., 2012), however, there is no direct evidence that PM H^+^-ATPase activation is involved in light-dependent nitrate uptake in leaves under nutrient sufficient condition. From our results, we speculate that energy production from photosynthesis activates PM H^+^-ATPase and thus nitrate uptake to maintain high nitrogen availability in leaves (**Figure 6**). The finding illustrates photosynthesis-dependent nitrate uptake in illuminated leaves is an interesting case of the carbon (C) nitrogen (N) interaction. Since molecular mechanism and physiological role of PM H^+^-ATPase activation in the context of C N interactions has never been demonstrated, our finding may provide a novel insight into the molecular mechanism of C N interaction in plant leaves.

## Materials and Methods

### Plant materials and growth conditions

*Arabidopsis thaliana* accession Columbia-0 (Col-0) was used as the wild-type plant.

Seeds were imbibed with tap water and stratified at 4°C for 3 to 4 days. Plants were grown in soil at 23°C under a 16-h light: 8-h dark cycle (6:00 h to 22:00 h light) with a photon flux of 50–100 μmol m^-2^ s^-1^. Soil was kept moistened and nutrition-rich by regular watering and fertilizer applications. Four- to five-week-old plants were transferred to a dark room for dark adaptation at 17:00 h to 18:00 h (ZT11 to ZT12) and leaves were harvested or treated at 11:00 h to 13:00 h (18 to 20 h after the start of dark adaptation). For white-light illumination in nitrate uptake assay, leaves of dark-adapted plants were harvested and leave the pieces of leaves under the same photon flux as growth condition for 1 h.

### Exogenous supplementation of carbon metabolites to leaves

Two fully expanded leaves (30–60 mg fresh weight) were harvested from independent dark-adapted plants and cut into two pieces, removing the main veins, upper edges, and lower edges. For sucrose analogue supplementation, pieces of leaves were floated on MilliQ water containing 30 mM mannitol (Nacalai), 30 mM sucrose (Nacalai), 30 mM turanose (Sigma), or 30 mM palatinose (Sigma) for 30 min.

For 2DG supplementation, pieces of leaves were floated on water containing (or not) 30 mM 2-deoxy glucose (Nacalai) for 1 h. Pieces of leaves were infiltrated in a syringe with water containing (or not) 60 mM mannitol, 30 mM sucrose, or 30 mM pyruvate-sodium (animal-free; Nacalai), and were floated on each solution for 30 min.

For the sucrose time-course experiment, pieces of leaves were infiltrated in a syringe with water containing 30 mM sucrose. Pieces of leaves were floated on the same sucrose solution and collected at the indicated times.

### Immunoblotting

Pieces of leaves were collected in tubes and flash-frozen in liquid nitrogen. Frozen leaf samples were homogenized using a pestle in solubilization buffer (2% [w/v] SDS, 1 mM EDTA, 20% [v/v] glycerol, 10 mM Tris-HCl [pH 6.8], 0.012% [w/v] CBB, 50 mM DTT, 1 mM PMSF, 2.5 mM NaF, 20 µM leupeptine [Leu]). Solubilized samples were centrifuged at 14,000 *×* g and 25°C for 5 min.

For MCPs sample, solubilization buffer was added to frozen MCPs. Identical volumes of supernatants were subjected to 9% sodium dodecyl sulfate-polyacrylamide gel electrophoresis. Immunoblotting was performed as described previously (Hayashi et al., 2010). Anti-H^+^-ATPase, anti-pThr, and anti-GFP (Roche; monoclonal) antibodies were used to evaluate the abundance of PM H^+^-ATPase, phosphorylated penultimate Thr of PM H^+^-ATPase (pThr), and GFP-fused SAUR30, respectively. Specificity of each antibodies were shown in Supplemental Figure S3.

Phosphorylation of PM H^+^-ATPase was determined based on band densities on anti-pThr and anti-PM H^+^-ATPase immunoblots using ImageJ software (version 1.53 f51).

### RNA extraction and cDNA synthesis

Total RNAs were extracted from leaves using NucleoSpin RNA Plant (MACHEREY-NAGEL) according to the manufacturer’s instructions. For RT**-**quantitative PCR, 400 ng of purified RNA were subjected to reverse transcription using the PrimeScriptRT Reagent Kit (TaKaRa).

### Preparation of complementary DNA libraries and RNA-seq

A TruSeq RNA Sample Prep Kit v. 2 (Illumina) were used for construction of complementary DNA libraries and the complementary DNA libraries were sequenced on a NextSeq 500 system (Illumina). NextSeq 500 pipeline software was used for base-calling of sequence reads. The reads used for mapping were selected by the length of 50 continuous nucleotides with quality values > 25. The selected reads were mapped to *Arabidopsis thaliana* transcripts using Bowtie software. Experiments in all conditions were repeated four times, independently; 11.3–22.2 million total reads per experiment were obtained. Gene-expression values are reported as transcripts per million (TPM) and log_2_ (fold change) values. Normalization of read counts and statistical analyses was performed using EdgeR software in the Degust v. 3.1.0 web tool, R software (v. 4.0.3), and R studio (v. 1.4.110). Values of FDR < 0.05 and log_2_(|fold change|) > 1 were used as cut-offs for selecting differentially expressed genes (DEGs). Reference transcriptome data were from KIN10-dependent genes (Baena-González et al., 2007).

### Construction of transient expression plasmids

The coding sequence (CDS) of *SAUR30* was cloned using Prime STAR MAX polymerase mix (TaKaRa) and the gene-specific primers shown in Supplemental Table S1. Amplified fragments were inserted into the pUC18 vector containing pro35S:sGFP:NOS terminator by in-fusion (TaKaRa). Before transfection, plasmids were purified from overnight-cultured competent *Escherichia coli* (DH5*α*) using a PureYield Plasmid Midiprep System (Promega) and the phenol-chloroform method.

### Preparation and transfection of MCPs

Enzymatic preparation of MCPs and PEG-Ca^2+^ transfection were conducted as described previously (Yoo *et al*., 2007) with minor modification. Briefly, mature leaves were harvested from 4–5-week-old plants and cell walls were enzymatically digested in a solution of Cellulase R-10 (Yakult) and Macerozyme R-10 (Yakult) for 2.5 h. Isolated MCPs (5.0 *×* 10^4^ protoplasts) in MMg (0.4 M mannitol, 4 mM MES-KOH [pH 5.7], 15 mM MgCl_2_) were transfected with the purified plasmids (20–30 μg) by adding an equal volume of PEG-Ca solution (40% PEG4000 [Sigma-Aldrich], 0.2 M mannitol, 100 mM CaCl_2_) for 5–6 min. After transfection, MCPs were washed three times and incubated in MKCa solution (0.4 M mannitol, 2 mM MES-KOH [pH 5.7], 20 mM KCl, 1 mM CaCl_2_) in 12-well plates for 4.5 h. Finally, MCPs were collected and stored at −80°C.

### RNA extraction and cDNA synthesis

Total RNA extraction and synthesis of complementary DNA (cDNA) from MCPs (approximately 400 MCPs) were performed using a SuperScript IV CellsDirect cDNA Synthesis Kit (Invitrogen) according to the manufacturer’s instructions.

### RT-PCR

RT-PCR detection of *SAUR30* and *UBQ5* expression was performed using ExTaq polymerase (TaKaRa) and the gene-specific primers listed in Supplemental Table S1.

### RT-quantitative PCR

Detection of *SAUR30* and *UBQ5* expression were performed using SYBR Green PCR Master Mix (Applied Biosystems), a StepOnePlus™ Real-time PCR System (Applied Biosystems), and the gene-specific primers listed in Supplemental Table S1.

### Stable-isotope-labelled nitrate-uptake assay

Six pieces of leaves from dark-adapted plants were washed with water and floated on water (dark and light treatment), 10 μM fusicoccin (Fc) in water, and 1 mM vanadate ammonium in Tris-HCl (pH 5.8). For vanadate treatment, pieces of leaves were incubated for 30 min in darkness and transferred to light for 1 h.

Treated leaves were transferred to 1 mM K^15^NO_3_ (atom% ^15^N: 99%) in water and incubated for 10 min, except for the no-K^15^NO_3_ control. Incubated leaves were washed twice in a large volume of water and collected into tubes. The samples were dried at 70°C for at least 20 h and the total N (% g dry weight [DW]) and ^15^N (atom%) contents were analyzed using a Flash2000-DELTAplus Advantage ConFlo? System (ThermoFisher Scientific) at Shoko Science Co., Ltd.

Influx of ^15^NO_3_ (µmol/gDW/h) was calculated from the total N (% gDW) and ^15^N (atom %) content, from which was subtracted the ^15^N content of the no-K^15^NO_3_ control.

### Data analysis and statistics

Plots were generated and statistical analyses were conducted using R studio with the ggplot2 and multcomp packages, respectively. ANOVA was conducted before a Tukey HSD or Dunnett’s *post hoc* test. The statistical results are listed in Supplemental Data File 2.

### Accession numbers

RNA-seq data reported are available in the DNA Data Bank of Japan (DDBJ) Sequenced Read Archive under the accession numbers DRA012961.

SAUR1, AT4G34770; SAUR3, AT4G34790; SAUR6, AT2G21210; SAUR7, AT2G21200; SAUR13, AT4G38825; SAUR14, AT4G38840; SAUR19, AT5G18010; SAUR22, AT5G18050; SAUR23, AT5G18060; SAUR24, AT5G18080; SAUR30, AT5G53590; SAUR31, AT4G00880; SAUR32, AT2G46690; SAUR38, AT2G24400; SAUR44, AT5G03310; SAUR50, AT4G34760; SAUR51, AT1G75580; SAUR52, AT1G75590; SAUR61, AT1G29420; SAUR65, AT1G29460; SAUR71, AT1G56150

## Data Availability

All data that support the findings of this study are included in the manuscript and Supplemental Data of this article. The raw data in this study are available from the corresponding author upon the request. RNA-seq raw data have been deposited in DNA Data Bank of Japan (DDBJ) Sequenced Read Archive under the accession numbers DRA012961.

## Acknowledgement

We thank Dr. Iris Finkemeier (University of Münster) for discussing on the metabolites flux in illuminated leaves, and Dr. Masaki Okumura (University of Minnesota) for providing the methodological advice. We are also grateful for the Drs. Koji Takahashi, Shin-ichiro Inoue, and Yuki Hayashi (Nagoya University) for providing the technical advice and generous supports.

## Author Contributions

S.N.K. and T.Kinoshita designed the research; S.N.K. and T.S. performed the research and analyzed data; All authors contributed to writing of the paper and reviewed the paper.

## References

Akiyama M, Sugimoto H, Inoue S, Takahashi Y, Hayashi M, Hayashi Y, Mizutani M, Ogawa T, Kinoshita D, Ando E, et al (2022) Type 2C protein phosphatase clade D family members dephosphorylate guard cell plasma membrane H^+^-ATPase. Plant Physiol 188: 2228–2240

Assmann SM, Simoncini L, Schroeder JI (1985) Blue light activates electrogenic ion pumping in guard cell protoplasts of Vicia faba. Nature 318: 285–287

Baena-González E, Rolland F, Thevelein JM, Sheen J (2007) A central integrator of transcription networks in plant stress and energy signalling. Nature 448: 938–942

de Bont L, Naim E, Arbelet-Bonnin D, Xia Q, Palm E, Meimoun P, Mancuso S, El-Maarouf-Bouteau H, Bouteau F (2019) Activation of plasma membrane H^+^-ATPases participates in dormancy alleviation in sunflower seeds. Plant Sci 280: 408–415

Chandran D, Reinders A, Ward JM (2003) Substrate specificity of the Arabidopsis thaliana sucrose transporter AtSUC2. J Biol Chem 278: 44320–44325

Cookson SJ, Williams LE, Miller AJ (2005) Light-dark changes in cytosolic nitrate pools depend on nitrate reductase activity in Arabidopsis leaf cells. Plant Physiol 138: 1097–1105

Delhon P, Gojon A, Tillard P, Passama L (1996) Diurnal regulation of NO3-uptake in soybean plants IV. Dependence on current photosynthesis and sugar availability to the roots. J Exp Bot 47: 893–900

DeWitt ND, Sussman MR (1995) Immunocytological localization of an epitope-tagged plasma membrane proton pump (H^+^-ATPase) in phloem companion cells. Plant Cell 7: 2053–2067

Evans JR, Clarke VC (2019) The nitrogen cost of photosynthesis. J Exp Bot 70: 7–15

Falhof J, Pedersen JT, Fuglsang AT, Palmgren M (2016) Plasma membrane H^+^-ATPase regulation in the center of plant physiology. Mol Plant 9: 323–337

Fernie AR, Roessner U, Geigenberger P (2001) The sucrose analog palatinose leads to a stimulation of sucrose degradation and starch synthesis when supplied to discs of growing potato tubers. Plant Physiol 125: 1967–1977

Hachiya T, Sakakibara H (2017) Interactions between nitrate and ammonium in their uptake, allocation, assimilation, and signaling in plants. J Exp Bot 68: 2501–2512

Harada A, Okazaki Y, Kinoshita T, Nagai R, Takagi S (2020) Role of proton motive force in photoinduction of cytoplasmic streaming in Vallisneria mesophyll cells. Plants 9: 376

Haruta M, Burch HL, Nelson RB, Barrett-Wilt G, Kline KG, Mohsin SB, Young JC, Otegui MS, Sussman MR (2010) Molecular characterization of mutant Arabidopsis plants with reduced plasma membrane proton pump activity. J Biol Chem 285: 17918–17929

Hayashi Y, Nakamura S, Takemiya A, Takahashi Y, Shimazaki K, Kinoshita T (2010) Biochemical characterization of in vitro phosphorylation and dephosphorylation of the plasma membrane H^+^-ATPase. Plant Cell Physiol 51: 1186–1196

Hoffmann RD, Portes MT, Olsen LI, Damineli DSC, Hayashi M, Nunes CO, Pedersen JT, Lima PT, Campos C, Feijó JA, et al (2020) Plasma membrane H^+^-ATPases sustain pollen tube growth and fertilization. Nature Commun 11: 2395

Inoue S, Kinoshita T (2017) Blue light regulation of stomatal opening and the plasma membrane H^+^-ATPase. Plant Physiol 174: 531–538

Kaiser WM, Huber SC (2001) Post-translational regulation of nitrate reductase: Mechanism, physiological relevance and environmental triggers. J Exp Bot 52: 1981–1989

Kinoshita T, Shimazaki K (1999) Blue light activates the plasma membrane H^+^-ATPase by phosphorylation of the C-terminus in stomatal guard cells. EMBO J 18: 5548–5558

Klein D, Stitt M (1998) Effects of 2-deoxyglucose on the expression of rbcS and the metabolism of Chenopodium rubrum cell-suspension cultures. Planta 205: 223–234

Li L, Sheen J (2016) Dynamic and diverse sugar signaling. Current Opinion in Plant Biology 33: 116–125

Matt P, Geiger M, Walch-Liu P, Engels C, Krapp A, Stitt M (2001) The immediate cause of the diurnal changes of nitrogen metabolism in leaves of nitrate-replete tobacco: A major imbalance between the rate of nitrate reduction and the rates of nitrate uptake and ammonium metabolism during the first part of the light period. Plant Cell Environ 24: 177–190

Minami A, Takahashi K, Inoue S, Tada Y, Kinoshita T (2019) Brassinosteroid induces phosphorylation of the plasma membrane H^+^-ATPase during hypocotyl elongation in Arabidopsis thaliana. Plant Cell Physiol 60: 935–944

Niittylä T, Fuglsang AT, Palmgren MG, Frommer WB, Schulze WX (2007) Temporal analysis of sucrose-induced phosphorylation changes in plasma membrane proteins of Arabidopsis. Mol Cell Proteom 6: 1711–1726

Nikolic M, Cesco S, Monte R, Tomasi N, Gottardi S, Zamboni A, Pinton R, Varanini Z (2012) Nitrate transport in cucumber leaves is an inducible process involving an increase in plasma membrane H^+^-ATPase activity and abundance. BMC Plant Biol 12: 2–13

Okumura M, Inoue S, Kuwata K, Kinoshita T (2016) Photosynthesis activates plasma membrane H^+^-ATPase via sugar accumulation. Plant Physiol 171: 580–589

Okumura M, Inoue S, Takahashi K, Ishizaki K, Kohchi T, Kinoshita T (2012a) Characterization of the plasma membrane H^+^-ATPase in the liverwort Marchantia polymorpha. Plant Physiol 159: 826–834

Okumura M, Takahashi K, Inoue S, Kinoshita T (2012b) Evolutionary appearance of the plasma membrane H^+^-ATPase containing a penultimate threonine in the bryophyte. Plant Sig Behav 7: 7–11

Pajak B, Siwiak E, Sołtyka M, Priebe A, Zieliński R, Fokt I, Ziemniak M, Jaśkiewicz A, Borowski R, Domoradzki T, et al (2020) 2-deoxy-D-glucose and its analogs: From diagnostic to therapeutic agents. Int J Mol Sci 21: 234

Palmgren MG (2001) Plant plasma membrane H^+^-ATPases: Powerhouses for nutrient uptake. Annu Rev Plant Physiol Plant Mol Biol 52: 817–845

Paponov IA, Paponov M, Teale W, Menges M, Chakrabortee S, Murray JAH, Palme K (2008) Comprehensive transcriptome analysis of auxin responses in Arabidopsis. Mol Plant 1: 321–337

Perchlik M, Tegeder M (2018) Leaf amino acid supply affects photosynthetic and plant nitrogen use efficiency under nitrogen stress. Plant Physiol 178: 174–188

Ren H, Park MY, Spartz AK, Wong JH, Gray WM (2018) A subset of plasma membrane-localized PP2C.D phosphatases negatively regulate SAUR-mediated cell expansion in Arabidopsis. PLoS Genet 14: e1007455

Robertson WR, Clark K, Young JC, Sussman MR (2004) An Arabidopsis thaliana plasma membrane proton pump is essential for pollen development. Genetics 168: 1677–1687

Sinha AK, Hofmann MG, Römer U, Köckenberger W, Elling L, Roitsch T (2002) Metabolizable and non-metabolizable sugars activate different signal transduction pathways in tomato. Plant Physiol 128: 1480–1489

Sondergaard TE, Schulz A, Palmgren MG (2004) Energization of transport processes in plants. Roles of the plasma membrane H^+^-ATPase. Plant Physiol 136: 2475–2482

Spartz AK, Ren H, Park MY, Grandt KN, Lee SH, Murphy AS, Sussman MR, Overvoorde PJ, Gray WM (2014) SAUR inhibition of PP2C-D phosphatases activates plasma membrane H^+^-ATPases to promote cell expansion in Arabidopsis. Plant Cell 26: 2129–2142

Sweetlove LJ, Beard KFM, Nunes-Nesi A, Fernie AR, Ratcliffe RG (2010) Not just a circle: Flux modes in the plant TCA cycle. Trends Plant Sci 15: 462–470

Szecowka M, Heise R, Tohge T, Nunes-Nesi A, Vosloh D, Huege J, Feil R, Lunn J, Nikoloski Z, Stitt M, et al (2013) Metabolic fluxes in an illuminated Arabidopsis rosette. Plant Cell 25: 694–714

Takahashi K, Hayashi K, Kinoshita T (2012) Auxin activates the plasma membrane H^+^-ATPase by phosphorylation during hypocotyl elongation in Arabidopsis. Plant Physiol 159: 632–641

Tong Y, Zhou JJ, Li Z, Miller AJ (2005) A two-component high-affinity nitrate uptake system in barley. Plant J 41: 442–450

Tränkner M, Tavakol E, Jákli B (2018) Functioning of potassium and magnesium in photosynthesis, photosynthate translocation and photoprotection. Physiol Plant 163: 414–431

Wong JH, Spartz AK, Park MY, D. M, Gray WM (2019) Mutation of a conserved motif of PP2C.D phosphatases confers SAUR immunity and constitutive activity. Plant Physiol 181: 353–366

Wong JH, Klejchová M, Snipes SA, Nagpal P, Bak G, Wang B, Dunlap S, Park MY, Kunkel E N, Trinidad B, Reed JW, Blatt MR, Gray WM (2021) SAUR proteins and PP2C.D phosphatases regulate H+-ATPases and K+ channels to control stomatal movements. Plant Physiol, 185(1), 256–273

Xiong Y, McCormack M, Li L, Hall Q, Xiang C, Sheen J (2013) Glucose-TOR signalling reprograms the transcriptome and activates meristems. Nature 496: 181–186

Zhang Y, Giese J, Kerbler SM, Siemiatkowska B, Perez de Souza L, Alpers J, Medeiros DB, Hincha DK, Daloso DM, Stitt M, et al (2021) Two mitochondrial phosphatases, PP2c63 and Sal2, are required for posttranslational regulation of the TCA cycle in Arabidopsis. Mol Plant 14: 1104–1118

