## Supplemental Information for "Photosynthetic-product-dependent Activation of Plasma Membrane H^+^-ATPase and Nitrate Uptake in Arabidopsis Leaves"

**Lists of Supplemental Data**

Supplemental materials and methods

Supplemental Figure S1. Illumination-induced PM $H^+$-ATPase phosphorylation.

Supplemental Figure S2. Overlapped DEGs in this study with reference KIN10-dependent DEGs.

Supplemental Figure S3. Specificity of antibodies used in this study.

Supplemental Figure S4. Regulation of PM $H^+$-ATPase phosphorylation by co-expression of SAUR30 and PP2C-Ds in mesophyll cell protoplasts.

Supplemental Data File 1. DEGs under light and sucrose treatment.

Supplemental Data File 2. Statistical results.

Supplemental Table S1. Primers used in this study.

#### Supplemental Materials and Methods

##### Construction of transient expression plasmids

The coding sequence (CDS) of *PP2C-D2* (AT3G17090), *PP2C-D6* (AT3G51370), and 3×FLAG were cloned using Prime STAR MAX polymerase mix (TaKaRa) and the gene-specific primers shown in Supplemental Table S1. Amplified fragments were inserted into the pUC18 vector containing pro35S:NOS terminator by in-fusion (TaKaRa). Before transfection, plasmids were purified from overnight-cultured competent *Escherichia coli* (DH5α) using a PureYield Plasmid Midiprep System (Promega) and the phenol-chloroform method.

##### Immunoblotting

Solubilization buffer was added to frozen MCPs. Identical volumes of supernatants were subjected to 9% sodium dodecyl sulfate-polyacrylamide gel electrophoresis.

Immunoblotting was performed as described previously (Hayashi et al., 2010). Anti-FLAG (Sigma; monoclonal) antibody was used to evaluate the abundance of FLAG-fused PP2C-Ds.

#### Supplemental Figure Legends

##### Fig S1. DEGs in this study overlapped with reference KIN10-dependent DEGs.

A, Phosphorylation of the penultimate Thr and abundance of PM $H^+$-ATPase in illuminated pieces of leaves using anti-pThr and anti-PM $H^+$-ATPase antibodies, respectively.

B, Phosphorylation of PM $H^+$-ATPase in illuminated pieces of leaves. Cross bars and error bars represent means and SD of three independent experiments.

##### Fig S2. DEGs in this study overlapped with reference KIN10-dependent DEGs.

A, Venn diagram of condition-specific and overlapping DEGs in light, sucrose-supplementation conditions, and the KIN10-dependent reference genes.

B, Heatmap of KIN10-dependent reference genes (62 genes) by log2 (fold-change) values. Ref.KIN10 values were obtained from expression changes in KIN10 overexpression (Baena-González *et al.*, 2007).

C, Schematic of KIN10-dependent global gene expression changes. KIN10 is the kinase subunit of the low-energy sensor SnRK1. Low energy activates SnRK1, induces reprogramming of global gene expression, and suppresses enzyme activities by phosphorylation. NR, nitrate reductase; SPS, sucrose-phosphate synthase; TFs, transcription factors.

##### Fig S3. Specificity of antibodies used in this study.

Entire immunoblotting results in Supplementary Figure S2 were shown here as the representative. Anti-pThr, anti-PM $H^+$-ATPase, and anti-GFP antibodies were used to evaluate the abundance of phosphorylated penultimate Thr of PM $H^+$-ATPase (pThr), PM $H^+$-ATPase, and GFP, respectively.

##### Fig S4. Regulation of PM $H^+$-ATPase phosphorylation by co-expression of SAUR30 and PP2C-Ds in mesophyll cell protoplasts.

Phosphorylation of the penultimate Thr and abundance of PM $H^+$-ATPase in MCPs. PP2C-D2 (M331K) and PP2C-D6 (M326K) lacks SAUR interaction (Wong *et al.*, 2019). The band that at 37–50 kDa is 3×FLAG-PP2C-D2 or 3×FLAG-PP2C-D6.

### Supplemental Figures

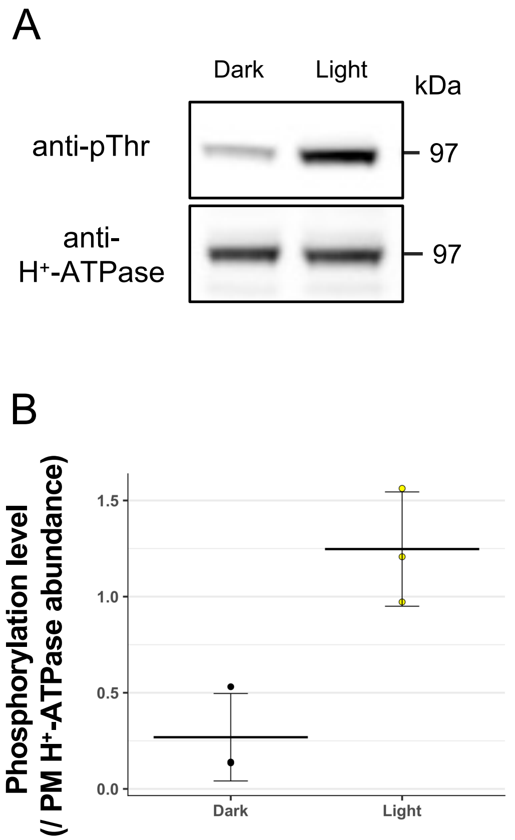


**Fig S1. Illumination-induced PM $H^+$-ATPase phosphorylation.**

A, Phosphorylation of the penultimate Thr and abundance of PM $H^+$-ATPase in illuminated pieces of leaves using anti-pThr and anti-PM $H^+$-ATPase antibodies, respectively.
B, Phosphorylation of PM $H^+$-ATPase in illuminated pieces of leaves. Cross bars and error bars represent means and SD of three independent experiments.

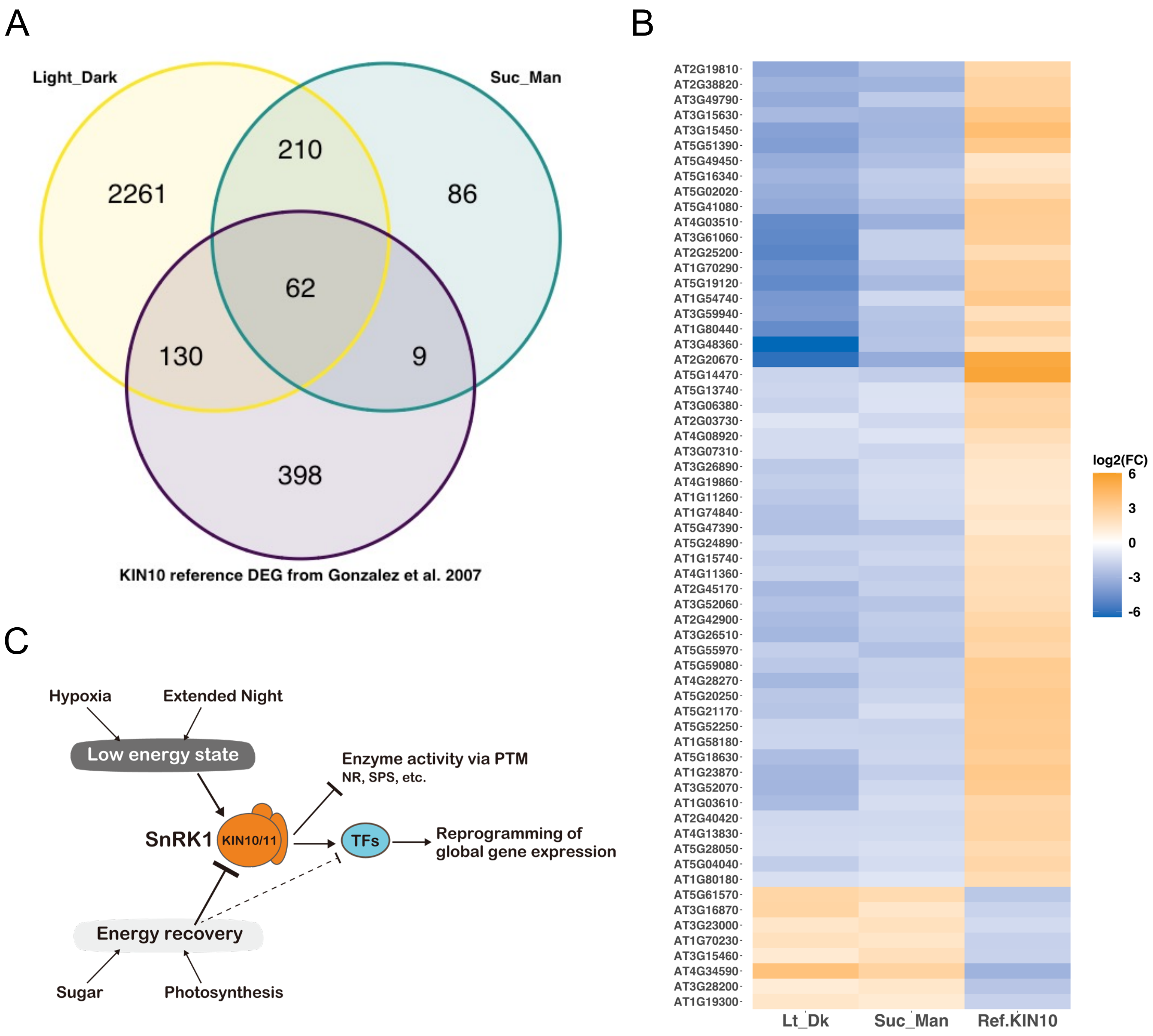


**Fig S2. Overlapped DEGs in this study with reference KIN10-dependent DEGs.**

A, Venn diagram of condition-specific and overlapping DEGs in light, sucrose-supplementation conditions, and the KIN10-dependent reference genes.

B, Heatmap of KIN10-dependent reference genes (62 genes) by log2 (fold-change) values. Ref.KIN10 values were obtained from expression changes in KIN10 overexpression (Baena-González *et al.*, 2007).

C, Schematic of KIN10-dependent global gene expression changes. KIN10 is the kinase subunit of the low-energy sensor SnRK1. Low energy activates SnRK1, induces reprogramming of global gene expression, and suppresses enzyme activities by phosphorylation. NR, nitrate reductase; SPS, sucrose-phosphate synthase; TFs, transcription factors.

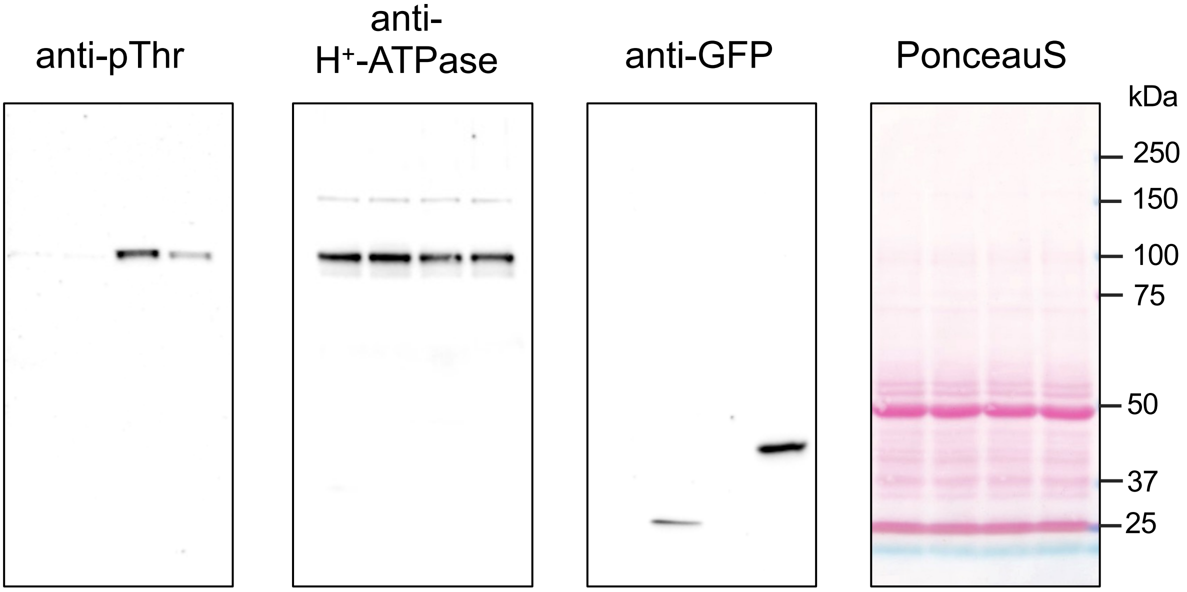


**Fig S3. Specificity of antibodies used in this study.**

Entire immunoblotting results in Supplementary Figure S2 were shown here as the representative. Anti-pThr, anti-PM $H^+$-ATPase, and anti-GFP antibodies were used to evaluate the abundance of phosphorylated penultimate Thr of PM $H^+$-ATPase (pThr), PM $H^+$-ATPase, and GFP, respectively.

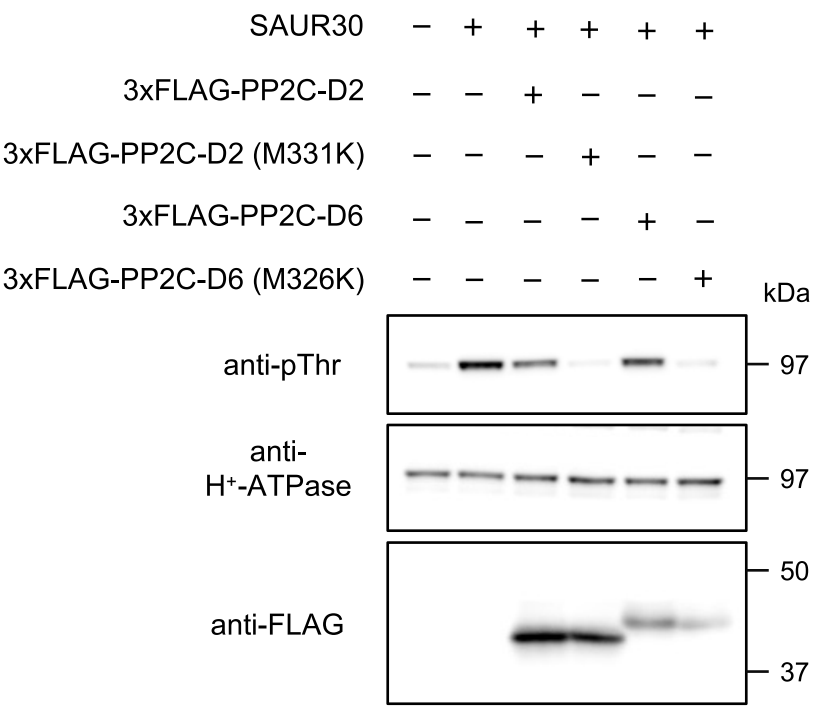


**Fig S4. Regulation of PM H$^+$-ATPase phosphorylation by co-expression of SAUR30 and PP2C-Ds in mesophyll cell protoplasts.**

Phosphorylation of the penultimate Thr and abundance of PM H$^+$-ATPase in MCPs. PP2C-D2 (M331K) and PP2C-D6 (M326K) lacks SAUR interaction (Wong *et al*., 2019). The band that at 37–50 kDa is 3×FLAG-PP2C-D2 or 3×FLAG-PP2C-D6.

**Supplemental Table S1. Primers used in this study.**

| Pimer for plasmid construction | |
|---|---|
| name | sequence |
| SalI-SAUR30_Fw | tatttacaattacagtcgacatgggttttgaag |
| SAUR30-lk-NcoI_Rv | gcccttgctcaccatgccagaaccgaaacatcgcaaatgg |
| NotI-SAUR30_Rv | ctgcagccgggcggccttagaaacatcgcaaatgg |
| SalI-FLAG_Fw | tatttacaattacagtcgacatggactacaaagac |
| linker-FLAG_Rv | gaaccaccaccacccttgtcatcgtcatccttg |
| linker-D6_Fw | ggtggtggtggttctatgttatcaacgt |
| NotI-D6_Rv_v2 | gatctgcagccgggcggccgcttagattttcttgg |
| linker-D2_Fw | ggtggtggtggttctatgtcaggttcattgatg |
| NotI-D2_Rv | gatctgcagccgggcggccgctcagtgttcaagagcac |
| D6_M326K_Fw | cgagagaagagatattctgatctaaag |
| D6_M326K_Rv | atatctcttctctcgtttctttgccgc |
| D2_M331K_Fw | cgtgagaagagatattcagatctaagga |
| D2_M331K_Rv | atatctcttctcacgtttcctcgctgc |

| Pimer for RT-PCR and RT-qPCR | |
|---|---|
| name | sequence |
| SAUR30_RT_q_Fw | TGGTGTTCAAGTTCCACTTCCA |
| SAUR30_RT_Rv | CGTGTCCCACCATAATCGCT |
| SAUR30_q_Rv | GTTATGGTTGTTGTGGCCGTG |
| UBQ5_Fw | CTTGAAGACGGCCGTACCCTC |
| UBQ5_Rv | CGCTGAACCTTTCAAGATCCATCG |
